# Toxoplasma activates host hypoxia inducible factor-1 by cytoplasmic trapping and lamp1-dependent lysosomal degradation of prolyl-hydroxylase 2

**DOI:** 10.1101/297333

**Authors:** Celia Florimond, Tongi Liu, Matthew Menendez, Kerstin Lippl, Christopher J. Schofield, Ira J. Blader

## Abstract

Hypoxia Inducible Factor-1 is a metazoan heterodimeric transcription factor that senses changes in O_2_ levels. HIF-1α subunit abundance is post-translationally regulated by prolyl-hydroxylase domain enzymes (PHDs), which use molecular O_2_ and α-ketoglutarate to hydroxylate two prolyl-residues in HIF-1α. Three PHDs have been identified and PHD2 is the most critical regulator of HIF-1α. HIF-1α can also be activated independently of hypoxia and in some cases this is due to changes in PHD2 abundance through poorly understood mechanisms. Previously, we reported that under O_2_-replete conditions that the intracellular parasite *Toxoplasma gondii* activates HIF-1 by reducing PHD2 protein abundance. Here, we demonstrate that *Toxoplasma* regulates PHD2 through a multistep process. First, PHD2 is a nucleocytoplasmic protein and *Toxoplasma* induces PHD2 cytoplasmic accumulation to separate it from nuclear HIF-1α. PHD2 is then degraded by lysosomes independently of the major autophagic processes, macroautophagy or chaperone-mediated autophagy. Rather, PHD2 interacts with the major lysosomal membrane protein, LAMP1, which is required for HIF-1 activation. These data therefore highlight for the first time that cytoplasmic trapping and subsequent lysosomal degradation of a host nucleocytoplasmic protein is a mechanism used by a microbial pathogen to regulate host gene expression.

## INTRODUCTION

Infections with the obligate intracellular parasite *Toxoplasma* lead to dramatic morphological and physiological changes to its host cells (Laliberte *et al.*, 2008, Blader *et al.*, 2014, Hakimi *et al.*, 2015). These include cytoskeletal rearrangements, alterations in membrane trafficking and changes in gene expression patterns due to differential activation of host transcription factors as well as epigenetic remodeling of the host genome. However, only a few of these changes are known to be required for parasite replication and thus far Hypoxia Inducible Factor-1 (HIF-1) and Interferon Regulatory Factor 3 are the only host transcription factors established as important for *Toxoplasma* growth (Spear *et al.*, 2006, Majumdar *et al.*, 2015).

HIF-1 is a transcription factor that is best known for its role in directing cellular responses to hypoxic stress, which it does by regulating the expression of genes involved in angiogenesis, glycolysis, apoptosis, cell proliferation and motility (Semenza, 2012). HIF-1 is a heterodimer composed of α and β subunits. In humans there are three HIF-α isoforms, which are associated with upregulation of different sets of genes, with HIF-1α and HIF-2α being more important in most biological processes (Greer *et al.*, 2012). HIF-1β protein is constitutively expressed while HIF-1α is normally undetectable at normoxia because it is rapidly degraded by the proteasome (Salceda *et al.*, 1997, Huang *et al.*, 1998). HIF-1α is only ubiquitylated and targeted to the proteasome after hydroxylation of two proline residues; an apparently irreversible reaction catalyzed by the Prolyl-Hydroxylase Domain (PHD) enzymes that use O_2_ and α-ketoglutarate as substrates and which react slowly with dioxygen (Bruick *et al.*, 2001, Epstein *et al.*, 2001). Hence, changes in O_2_ availability determine HIF-α protein levels and the activity of HIF. There are three known HIF-α modifying PHDs (PHD1-3) with PHD2 being the critical negative regulator of HIF-1α (Berra *et al.*, 2003, Appelhoff *et al.*, 2004). While much progress has been made in understanding how PHD2 interacts with and regulates HIF-1α, little is known about PHD2 turnover or localization although overexpression-based assays suggest that PHD2 is a dynamic nucleocytoplasmic protein that contains a nuclear export signal (NES) at its N-terminus and a non-canonical nuclear localization signal (NLS) in the middle of the protein adjacent to its catalytic domain (Steinhoff *et al.*, 2009, Yasumoto *et al.*, 2009, Pientka *et al.*, 2012).

The goal of this study was to determine how *Toxoplasma* regulates PHD2. Here, we report that *Toxoplasma* induces cytoplasmic accumulation of PHD2, which separates it from its substrate, HIF-1α, in the nucleus. In the cytoplasm, PHD2 associate with and is by degraded by host lysosomes. We further demonstrate that PHD2 interacts with the major lysosomal membrane protein, LAMP1, and that loss of LAMP1 leads to reduced HIF-1 activation and altered nucleocytoplasmic trafficking of PHD2. These data therefore highlight for the first time that cytoplasmic trapping of a host nucleocytoplasmic protein is a mechanism used by a microbial pathogen to regulate host gene expression.

## RESULTS

### Toxoplasma Induces Cytoplasmic Accumulation of PHD2

Previously, we demonstrated that under O_2_-replete conditions *Toxoplasma* infections activates HIF-1 by decreasing HIF-1α prolyl hydroxylation and that this was accompanied by a concomitant decrease in PHD2 protein levels. To directly compare PHD activity in *Toxoplasma*-infected cells, cell extracts from mock or parasite-infected cells were incubated with recombinant GST-tagged HIF-1αODD (the domain in HIF-1α containing the prolines modified by PHD2) immobilized on GST-beads (Tuckerman *et al.*, 2004). After extensive washing, HA-tagged VHL protein, which specifically binds prolyl hydroxylated HIF-1α, was added to the beads. The beads were then extensively washed and bound VHL identified by Western blotting. Unexpectedly, a significant increase in VHL levels were observed in samples incubated with lysates from the *Toxoplasma*-infected cells (Figure 1A). This was not due to the *Toxoplasma* prolyl hydroxylases TgPhyA or TgPhyB (Xu *et al.*, 2012) modifying HIF-1α since neither enzymes could catalyze HIF-1α hydroxylation (Supplemental Figure 1). Increased PHD activity was accompanied by an increase of PHD2 protein in the cell extracts (Figure 1B), which was surprising since PHD2 protein levels are decreased in *Toxoplasma*-infected cells (Wiley *et al.*, 2010).

**Figure 1:**
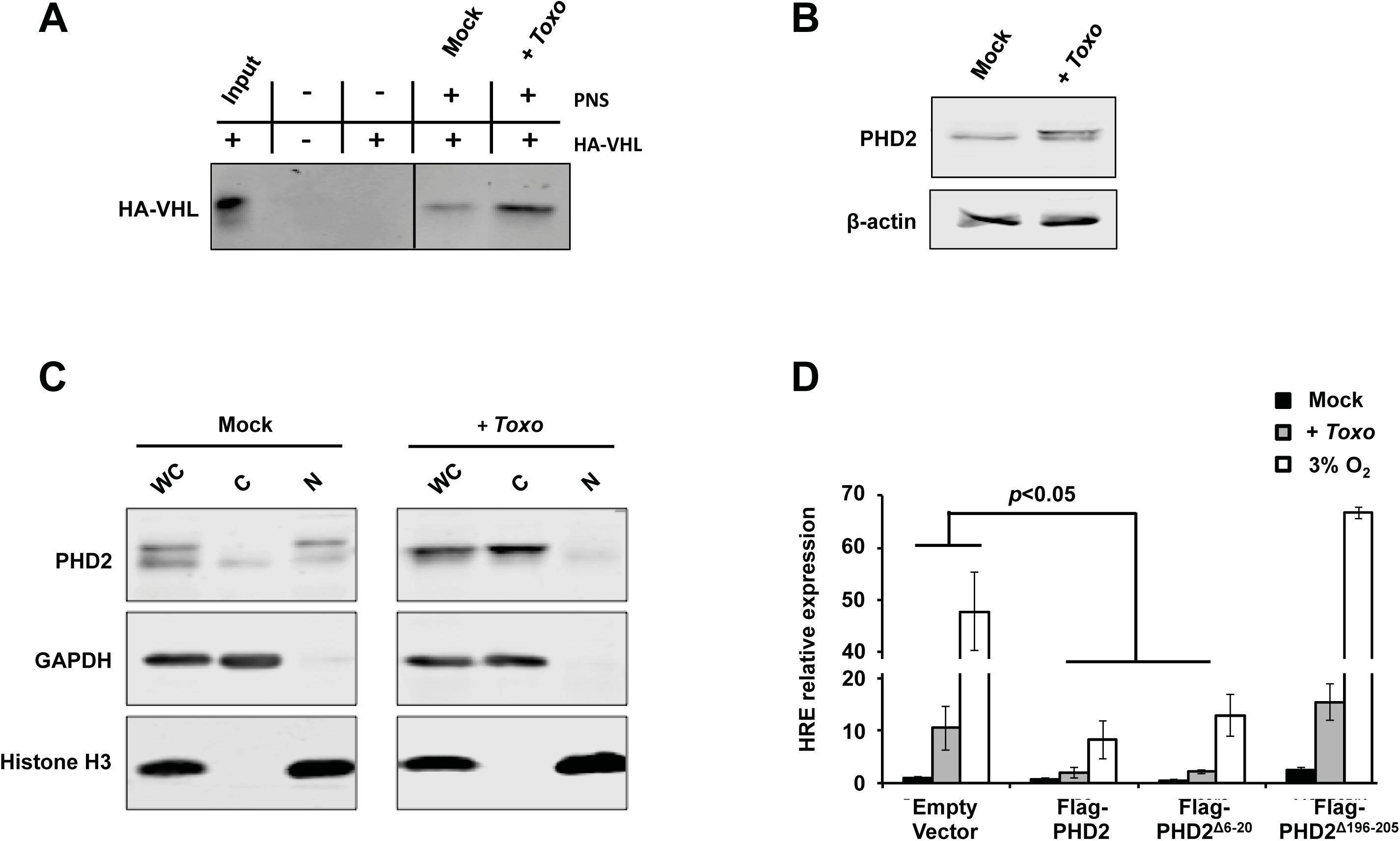
PHD2 Cytoplasmic Accumulation is Necessary for HIF-1 Activation in *Toxoplasma*-Infected Cells. **(A)** GST-HIF-1α bound to glutathione-agarose beads was incubated in the absence of presence of post-nuclear supernatant (PNS) extracts from mock or *Toxoplasma*-infected HFFs and recombinant min HA-VHL. The reactions were analyzed by Western blotting with anti-HA antibody. **(B)** Western-blot analysis of PHD2 abundance of the PNS extracts used for the VHL capture assay. **(C)** Western-Blot analysis of nuclear and cytoplasmic extracts from mock or parasite-infected HFFs. **(D)** Cells co-transfected with the pHRE-luc luciferase reporter and either wild-type PHD2 or PHD2 NES (PHD2∆^6-20^) or NLS (PHD2∆^196-205^) mutants were infected with parasites. After 18 h, cells were harvested and luciferase activity measured. Shown are the means and standard deviations of three independent assays.

The cell extracts used for the VHL capture assay were generated from post-nuclear supernatants. Since PHD2 is a nucleocytoplasmic protein (Yasumoto *et al.*, 2009, Pientka *et al.*, 2012) and PHD2 has been proposed to predominantly hydroxylate HIF-1α in the nucleus (Pientka *et al.*, 2012), we hypothesized that increased PHD2 levels observed in the cell extracts was due to *Toxoplasma* altering PHD2 nucleocytoplasmic trafficking. To test this, we compared PHD2 protein levels in nuclear and cytosolic extracts prepared from mock- and parasite-infected cells. Consistent with data from the VHL capture assays PHD2 protein accumulated in the cytoplasm of *Toxoplasma*-infected cells (Figure 1C). We next tested whether nucleocytoplasmic trafficking of PHD2 was important for HIF-1 activation by transfecting cells with a HIF-1 luciferase reporter and plasmids expressing either wild-type PHD2 or PHD2 mutants lacking either nuclear export or localization signals (Yasumoto *et al.*, 2009, Pientka *et al.*, 2012). HIF-1 regulated luciferase activity was reduced in parasite-infected cells expressing either wild-type PHD2 or the PHD2 mutant lacking the nuclear export signal. By contrast, HIF-1-regulated luciferase activity was unaltered in parasite-infected cells transfected with the PHD2 lacking its nuclear localization signal (NLS) (Figure 1D). In contrast, hypoxic activation of HIF-1 was unaffected in cells expressing the NLS-lacking PHD2 mutant indicating that this mutant had no unexpected off target effects on the HIF-1 pathway. It was unclear why the NLS mutant would have a dominant negative effect and inhibit endogenous PHD2, but may reflect an ability for PHD2 to form complexes including multimers in the cytoplasm (McDonough *et al.*, 2006, Lee *et al.*, 2016). Taken together, these data indicate that *Toxoplasma* activation of HIF-1 requires cytoplasmic accumulation of PHD2, which most likely acts to separate the enzyme from its substrate within the nucleus.

### PHD2 is Degraded By Host Lysosomes

Having established that *Toxoplasma* induces cytoplasmic accumulation of PHD2 to activate HIF-1, we sought to assess the localization and fate of PHD2 within the cytoplasm. Because PHD2 was reported to associate with membranes (Barth *et al.*, 2009), we assessed PHD2 levels in soluble (S100) and membrane (P100) fractions prepared from post nuclear extracts from mock and parasite-infected cells. Infection led to a significant increase in the amount of membrane-associated PHD2 (Figure 2A). We next examined the nature of the interaction between PHD2 and membranes by sodium carbonate extraction, which revealed that a small amount of membrane-associated PHD2 behaved as an integral membrane protein perhaps due to a post translational modification such as lipidation that results in membrane anchoring (Figure 2B). Finally, we assessed the topology of membrane-associated PHD2 by determining whether it was protected from protease treatment. We found that a significant fraction of membrane-associated PHD2 from both uninfected and infected cells was protected from Proteinase K degradation indicating its luminal localization (Figure 2C).

**Figure 2:**
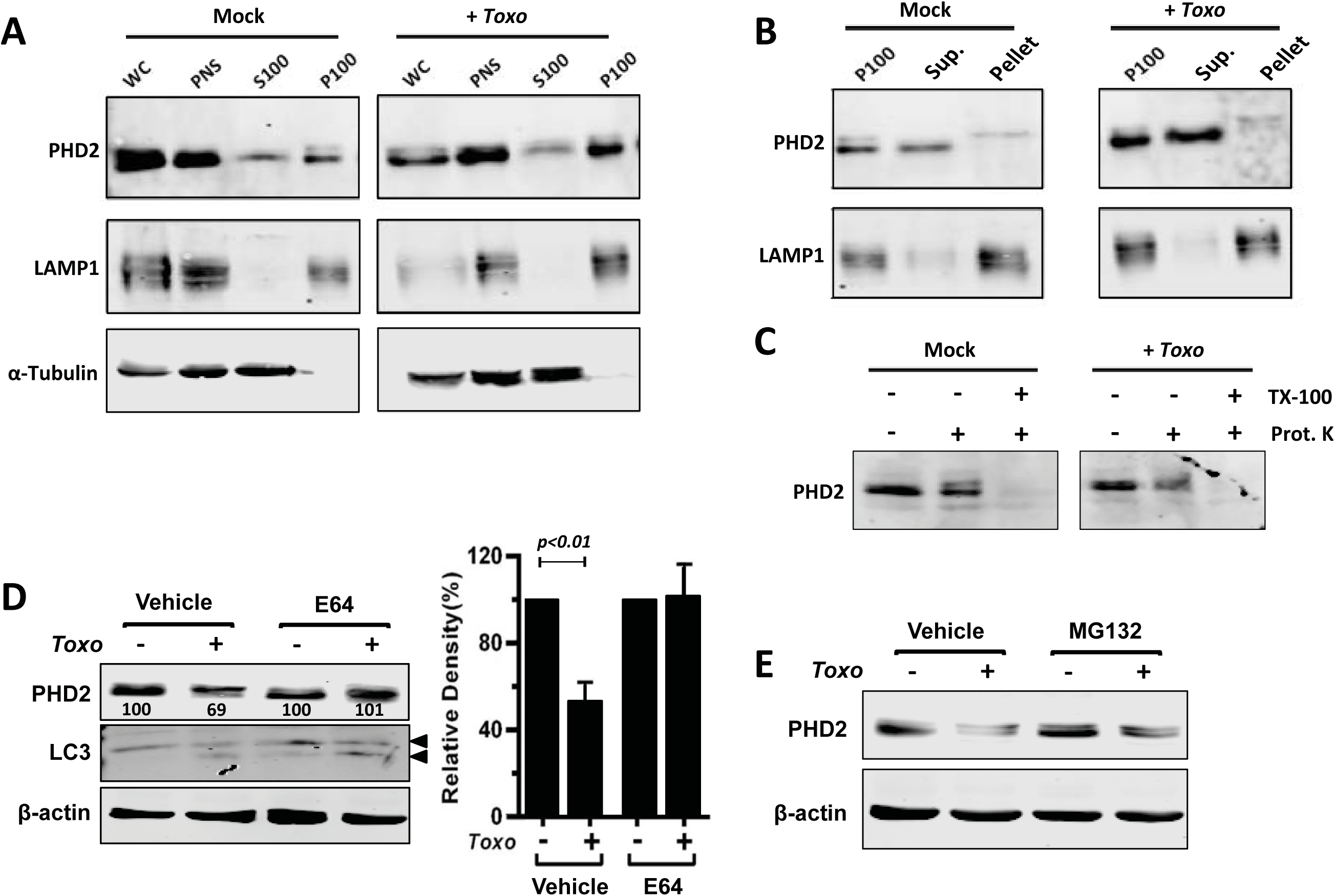
*Toxoplasma* Stimulates Lysosomal Degradation of PHD2. **(A)** Equivalent cell volumes of whole extracts, PNS, S100, and P100 fractions from mock- or parasite-infected cells (18 hpi) were Western Blotted to detect PHD2 and LAMP1. **(B)** PHD2 was detected in P100 fractions from mock or parasite-infected cells were incubated in the absence or presence of carbonate extraction buffer. **(C)** P100 fractions from mock or parasite-infected cells were incubated in the absence or presence of Proteinase K. In addition, Triton X-100 was added to Proteinase K treated samples to disrupt all membranes. **(D)** PHD2, LC3 (a positive control for E64 which accumulates lipidated-LC3 (bottom arrow) (Tanida *et al.*, 2004)), and β-actin as loading were detected in lysates from mock or parasite-infected cells treated with or without E64 (**E)** HFFs were mock or parasite-infected for 18 h and treated with or without MG132 for the last 6 h. Whole cell lysates were prepared and probed to detect PHD2 and β-actin.

Our findings that PHD2 is lumenally localized and its abundance is down regulated in *Toxoplasma*-infected cells suggested that infection induced PHD2 degradation within lysosomes. To test this, we compared PHD2 protein levels between mock- or *Toxoplasma*-infected cells incubated in the absence or presence of the thiol lysosomal protease inhibitor, E64. We found that PHD2 levels remained high in E64-treated parasite-infected cells (Figure 2D). In contrast, the proteasome inhibitor, MG132, had no effect on endogenous PHD2 levels in parasite-infected cells (Figure 2E). Similarly, down regulation of epitope-tagged PHD2 ectopically expressed in MEFs was resistant to MG132 while the inhibitor did increase the NF-κB regulating protein, IκBalpha, whose degradation was previously reported to be sensitive to MG132 (Palombella *et al.*, 1994) (Supplemental Figure 2).

Given that E64 reversed the effects of infection on PHD2 protein levels, we sought to examine the association of PHD2 with lysosomes. Our initial attempts to assess PHD2 localization by immunofluorescence assays were inconclusive because cytoplasmic PHD2 interfered with imaging lysosome-associated PHD2 (not shown). Therefore, post nuclear supernatant extracts from uninfected cells were fractionated by density gradient centrifugation on a continuous Percoll gradient to biochemically assesses PHD2 localization (Figure 3A). As previously reported (Barth *et al.*, 2009), PHD2 was detected in a wide range of fractions although it was most highly abundant in fractions containing lysosomal markers. We also noted that the slower migrating PHD2 species that pelleted following carbonate extraction (Figure 2E) was enriched in the lysosomal fractions. Similar results were noted using an Optiprep step flotation gradient (Figure 3B).

**Figure 3:**
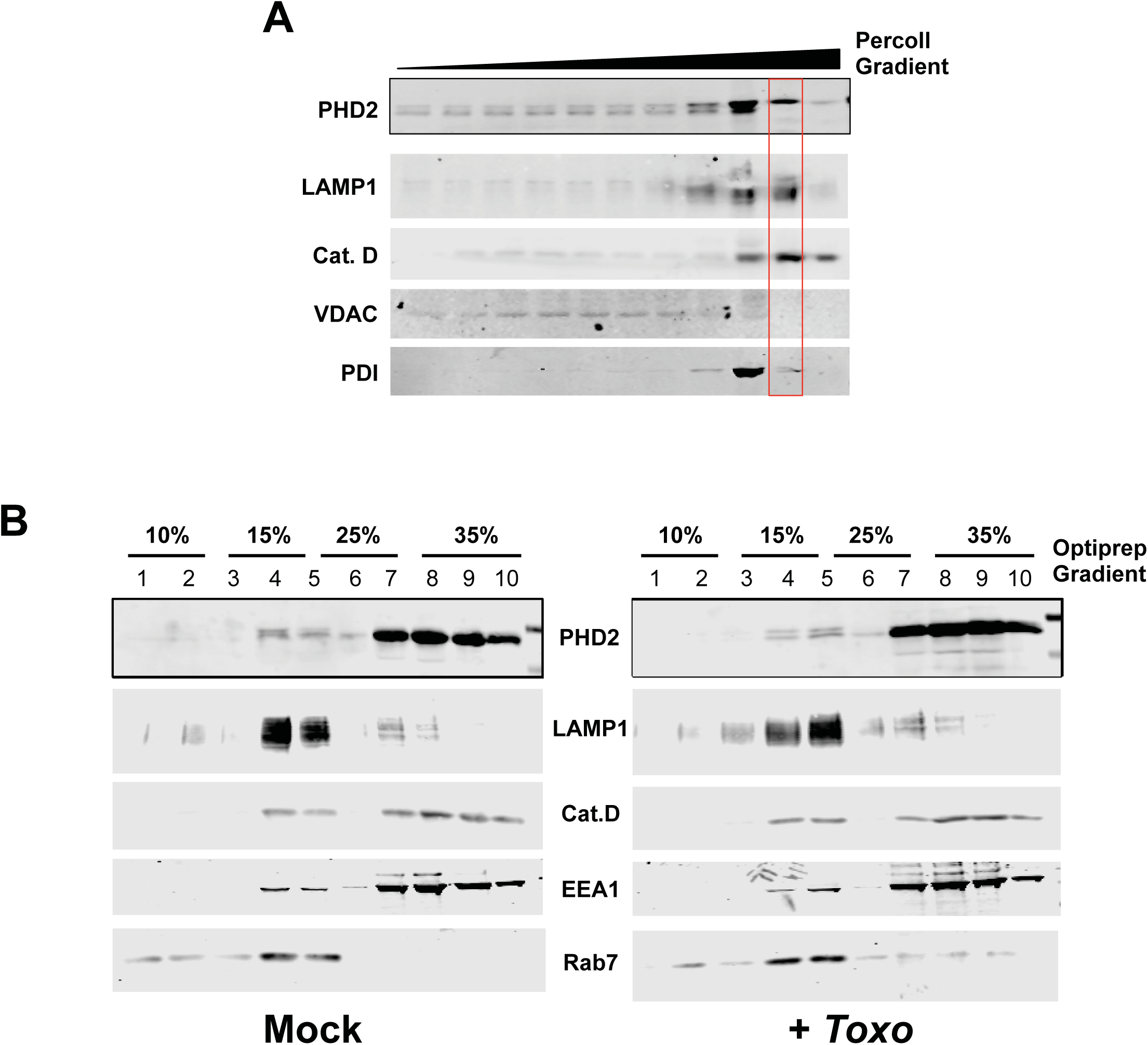
(A) Cell fractionation on Percoll gradient performed of post-nuclear extract from HFF Mock condition. (B) Cell fractionation and flotation assay on Optiprep step gradient of post-nuclear extracts from mock or infected HFF; LAMP1 was used as a membrane marker for Endosomes/Lysosomes; Cathepsin D as a lumen marker of Lysosomes; EEA1 as a marker for Early endosome membrane; VPS4 as a marker for ESCRT system.

### PHD2 Interacts with LAMP1

Two major autophagic pathways target cytosolic proteins for lysosomal degradation. First, classical autophagy (macro- and microautophagy) is dependent on the autophagic regulatory protein, ATG5 (Mizushima *et al.*, 1998, Mizushima *et al.*, 2001). However, PHD2 protein levels were similarly decreased in mock- or parasite-infected cells transfected with ATG5 or negative control siRNAs (Figure 4A). In addition, a large-scale RNAi screen failed to identify other host autophagy proteins and regulators as important for *Toxoplasma* growth under either normoxic or hypoxic conditions (Menendez *et al.*, 2015) suggesting that they are dispensable for HIF-1α activation in *Toxoplasma*-infected cells. Chaperone-mediated autophagy is a second lysosomal degradative pathway for cytoplasmic proteins and requires the lysosomal membrane protein LAMP2A (Cuervo *et al.*, 1996). Western blotting lysates from mock and parasite-infected cells using a LAMP2A-specific antibody revealed that *Toxoplasma*-infection led to significantly reduced LAMP2A protein levels (Figure 4B), which would impede chaperone-mediated autophagy activity (Cuervo *et al.*, 2000b, Cuervo *et al.*, 2000a). However, total LAMP2 levels were increased in parasite-infected cells indicating that decreased LAMP2A protein levels most likely occur due to post-transcriptional events since LAMP2 isoforms are synthesized by alternative splicing of the same transcript (Cuervo *et al.*, 2000b).

**Figure 4:**
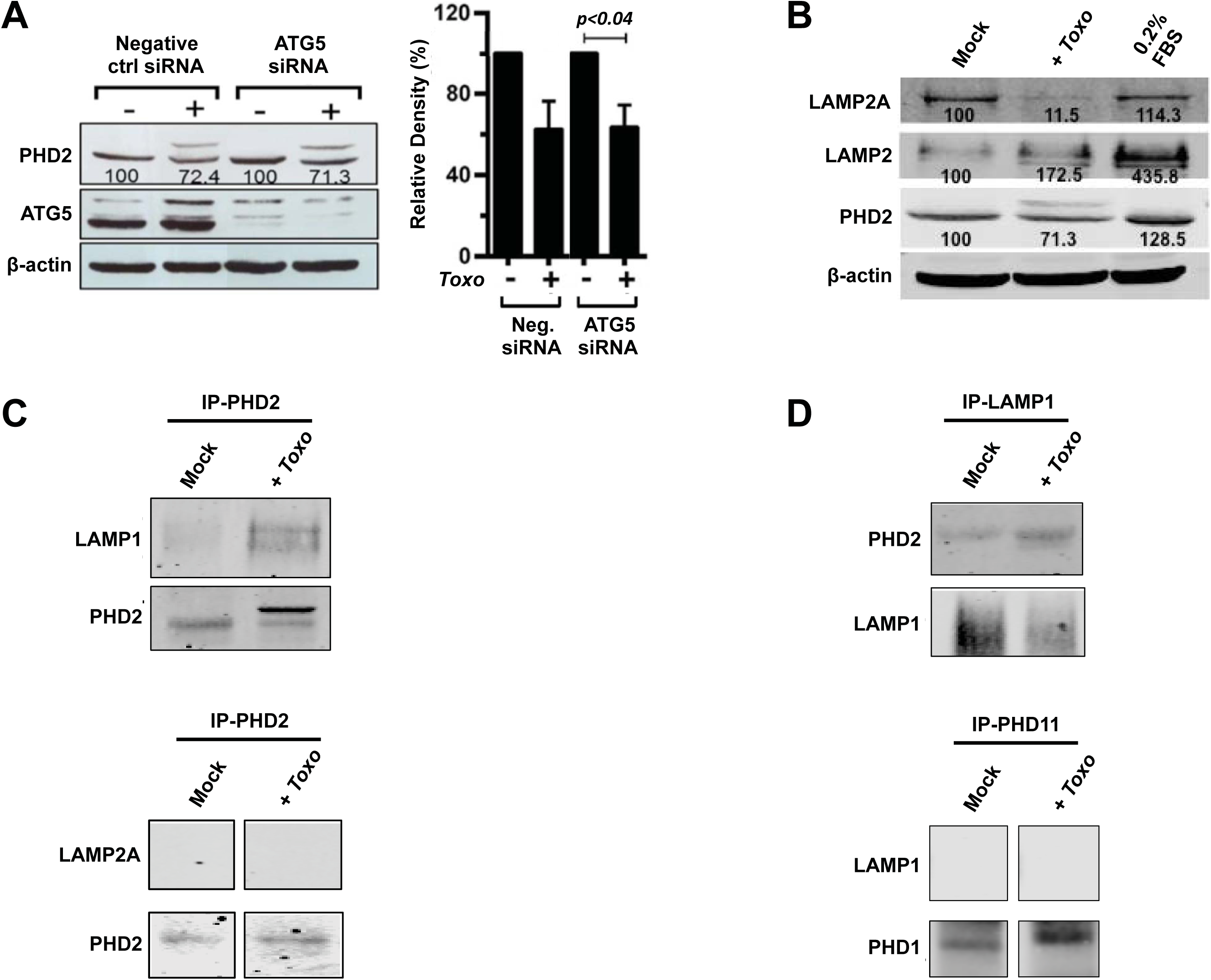
PHD2 Interacts with LAMP1. **(A)** HeLa cells were transfected with negative control or ATG5 siRNAs and 48 h later mock or parasite-infected for 18 h. Lysates were prepared and Western blotted to detect PHD2, ATG5, or β-actin. **(B)** Lysates prepared mock- and parasite-infected cells (18 hpi) or serum-starved cells (0.2% FBS) for 18h were Western blotted to detect LAMP2A, total LAMP2, PHD2, and β-actin. **(C)** PHD2 immunoprecipitates from mock or parasite-infected HFFs were Western blotted to detect LAMP1 and LAMP2A. **(D)** LAMP1 immunoprecipitates from mock or parasite-infected HFFs were Western blotted to detect PHD2 and PHD1.

Although reduced LAMP2A protein levels suggested that chaperone-mediated autophagy was not involved in regulating PHD2 in *Toxoplasma*-infected cells, we tested whether any remaining LAMP2A could interact with PHD2. However, LAMP2A was not detected in PHD2 immunoprecipitates from either mock- or parasite-infected cells (Figure 4C). In parallel, PHD2 immunoprecipitates were Western blotted to detect the major lysosomal membrane protein, LAMP1. In contrast to LAMP2A, LAMP1 co-immunoprecipitated with PHD2 and the PHD2/LAMP1 interaction increased in *Toxoplasma*-infected cells (Figure 4D). PHD2 similarly was detected in LAMP1 immunoprecipitates. The interaction between LAMP1 and PHD2 appeared specific since LAMP1 could not be detected when another HIF hydroxylase PHD1 was immunoprecipitated.

### LAMP1 Is Required for HIF-1 Activation in Toxoplasma-Infected Cells

LAMP1 is the most abundant lysosomal membrane protein (Chen *et al.*, 1985) and despite its discovery over 30 years ago defining its cellular function has remained elusive since LAMP2 can compensate for loss of LAMP1 (Andrejewski *et al.*, 1999, Huynh *et al.*, 2007). Given that PHD2 degradation occurred in lysosomes, we first sought to examine whether PHD2 localization to lysosomes were altered in LAMP1-deficient cells. But despite repeated attempts using a variety of protocols (and in contrast to other cells that we used) we were unable to purify intact lysosomes (data not shown) from either wild-type or LAMP1KO murine embryonic fibroblasts (MEFs). As an alternative, we examined whether interactions between LAMP1 and PHD2 were required for HIF-1 activation. In Figure 1, we showed that PHD2 nuclear export was required for HIF-1 activation in *Toxoplasma*-infected cells. Therefore, we first compared PHD2 nucleocytoplasmic distribution in mock- or parasite-infected WT or LAMP1KO MEFs. In contrast to the human fibroblasts used in Figure 1, MEFs had higher steady-state levels of cytoplasmic PHD2 (Figure 5A) than observed suggesting species and/or cell specific differences in PHD2 nucleocytoplasmic distribution. *Toxoplasma* infection led to significantly reduced levels of nuclear PHD2 in wild-type MEFs. In contrast, PHD2 levels in nuclei of *Toxoplasma*-infected LAMP1KO cells were not decreased.

**Figure 5:**
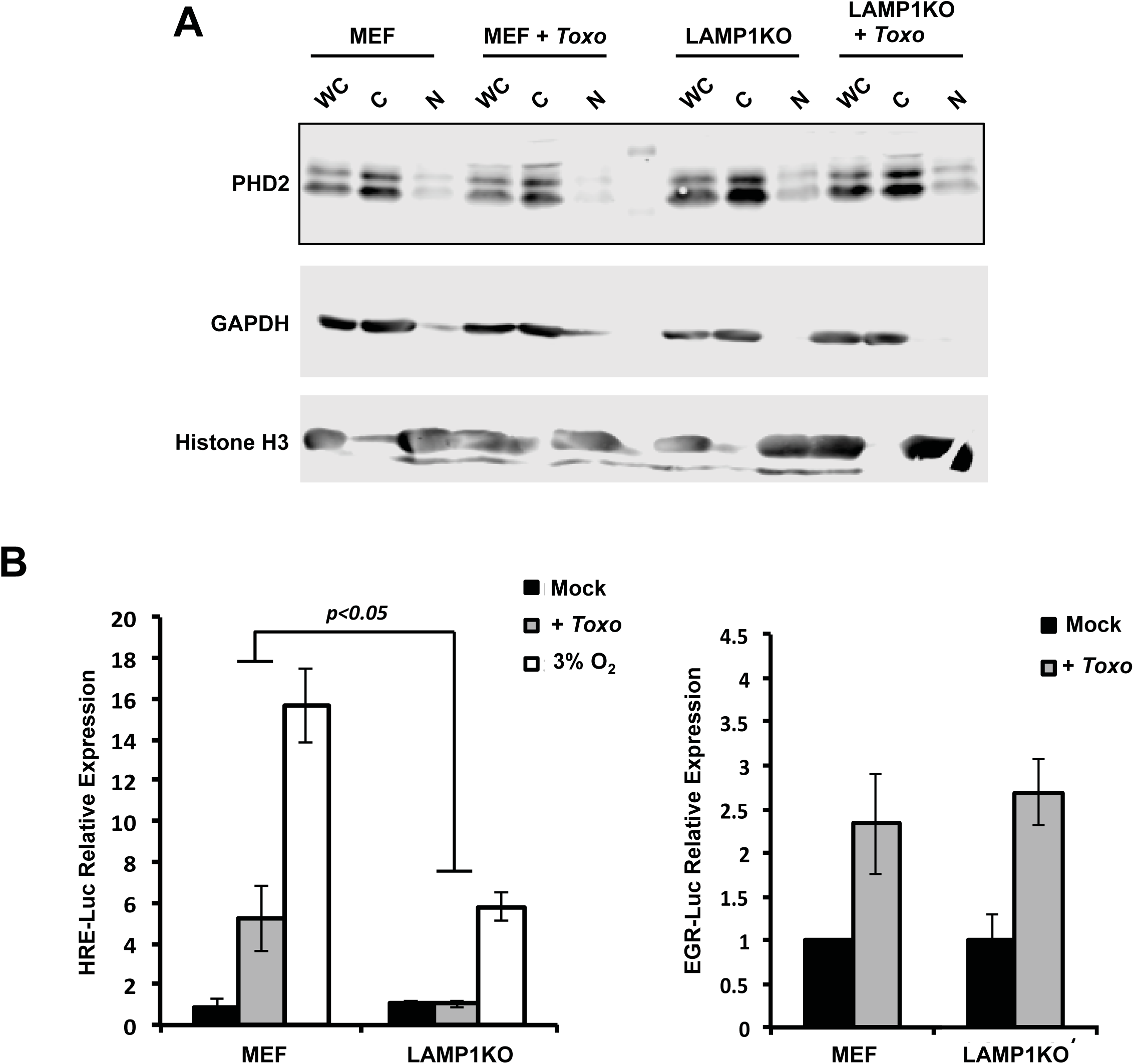
LAMP1 is Required for Cytoplasmic Accumulation of HIF-1α and HIF-1 Activation. (A) Cell fractionation and Western-Blot analysis of MEF and Lamp1^-/-^ cells in mock or infected conditions; GAPDH is used as a cytoplasmic marker and Histone H3 as a nuclear marker. (C) and (D) MEF and LAMP1^-/-^ cells were infected with *Toxoplasma*. The activation of HIF1 was defined by Luciferase expression under the control of HRE (C); or under the control of EGR (D).

Next, we determined what effect loss of LAMP1 had *Toxoplasma* activation of HIF-1. Thus, wild-type and LAMP1KO MEFs were transfected with a HIF-1-luciferase reporter and then infected for 18 h at which time luciferase activity was measured. The data indicated that HIF-1 activation was abrogated in *Toxoplasma*-infected LAMP1-deficient cells (Figure 5B). An inability to activate HIF-1 was not a general defect in host-parasite signaling since activation of the host EGR2 transcription factor was similar in wild type and LAMP1KO cells (Figure 5C). Finally, we assessed whether loss of LAMP1 impacted HIF-1 activation at 3% O_2_, which represents tissue oxygen levels that are not considered hypoxia. We found that HIF-1 activation by exposure of cells to 3% O_2_ was reduced in the LAMP1KO cells most likely due to the constitutively elevated PHD2 protein levels in the LAMP1KO cells (Figure 5B),

## DISCUSSION

Intracellular pathogens employ diverse strategies to establish replicative niches within their host cells. These include altering host cell transcription, membrane trafficking, cytoskeletal elements, protein and mRNA stability, and signaling. In our previous work, we demonstrated that *Toxoplasma* activates HIF-1 by inhibiting PHD2 activity (Wiley *et al.*, 2010) and the goal of this work was to define how the parasite regulates PHD2. Our serendipitous observation that cytoplasmic extracts from *Toxoplasma*-infected cells contained high levels of PHD2 protein as well HIF-1-directed prolyl hydroxylase activity led to our discovery that *Toxoplasma* activates HIF-1 by inducing cytoplasmic trapping of PHD2. Within the cytoplasm, PHD2 is targeted to lysosomes and degraded. These data represent, to our knowledge, the first example of such a degradation mechanism deployed by a microbial pathogen to activate a host transcription factor.

Degradation of cytosolic proteins within lysosomes is largely accomplished through either ATG5-dependent autophagy or chaperone-mediated autophagy. While ATG5-dependent autophagic processes remain active in *Toxoplasma*-infected cells (Wang *et al.*, 2009, Khaminets *et al.*, 2010, Selleck *et al.*, 2013, Choi *et al.*, 2014), chaperone-mediated autophagy had not been examined. Our observations that LAMP2A levels are significantly decreased in parasite-infected cells, however, would suggest that chaperone-mediated autophagy is generally reduced since LAMP2A is required for chaperone-mediated autophagy (Cuervo *et al.*, 2000b, Cuervo *et al.*, 2000a). The implications for this finding are not yet clear but could represent another approach to facilitate *Toxoplasma* growth since HIF-1α has been reported to be a chaperone-mediated autophagy substrate (Hubbi *et al.*, 2013). GAPDH is another chaperone-mediated autophagy substrate whose dysregulation in *Toxoplasma*-infected cells may facilitate parasite growth since *Toxoplasma* increases host cell glycolysis (Menendez *et al.*, 2015). Proteins that are substrates for chaperone-mediated autophagy contain a KFERQ motif that associates with cytoplasmic hsc70 (Cuervo *et al.*, 1994). More recently it was noted that the KFERQ motifs could also direct proteins to lysosomes via a second pathway, endosomal microautophagy, which differs from chaperone-mediated autophagy because is not dependent on LAMP2A (Sahu *et al.*, 2011). We do not believe that PHD2 is a substrate for endosomal microautophagy since PHD2 does not possess a canonical KFERQ motif. However, we are able to co-IP PHD2 with hsc70 (not shown), which is required for both endosomal and chaperone-mediated autophagy (Kaushik *et al.*, 2012), suggesting that lysosomal targeting of PHD2 occurs through a novel pathway.

Although PHDs are often thought to modify HIF-1α in the cytoplasm, data supporting this model were largely based on earlier data that HIF-1α accumulated in nuclei of hypoxic cells and that upon reoxygenation HIF-1α would redistribute to the cytoplasm (Kallio *et al.*, 1998). However, more recent work revealed that PHD2 substantially regulates HIF-1α in the nucleus (Pientka *et al.*, 2012). While both of these studies were based on overexpression of either HIF-1α or PHD2, our work examined endogenous PHD2 levels and our conclusions support those of (Pientka *et al.*, 2012). However, neither our work nor that described in (Pientka *et al.*, 2012) have defined how PHD2 nucleocytoplasmic trafficking is regulated. This is difficult to address in *Toxoplasma*-infected cells since PHD2 nuclear export is CRM1-dependent but the inhibitor of this process, leptomycin B, possesses anti-parasitic activity (our unpublished data and (Francia, 2013)). However, the observed PHD2 accumulation in nuclei of the wild-type, but not LAMP1KO murine MEFs, support a model in which *Toxoplasma* promotes the nuclear export of PHD2 to the cytoplasm where it engages LAMP1 to facilitate its lysosomal degradation. Beyond our *Toxoplasma* studies, this mechanism may impact other diseases and their therapies. For example, the PHDs are current therapeutic targets for anemia and other ischemia related diseases with inhibitors for them in late stage clinical trials and the results presented here suggest a new way of regulating PHD2 activity by modulating its interaction with LAMP1 (Chan *et al.*, 2016, Maxwell *et al.*, 2016, Haase, 2017, Martin *et al.*, 2017).

LAMP1 was discovered over 30 years and is the major lysosomal membrane protein (Chen *et al.*, 1985). Deletion of LAMP1 has no dramatic effect on lysosomal function, which is likely due to increased expression of LAMP2 that appears to be functionally redundant with LAMP1 (Andrejewski *et al.*, 1999, Eskelinen *et al.*, 2004, Huynh *et al.*, 2007). LAMP2 can function independently of LAMP1 in chaperone-mediated autophagy and in lysosomal plasma membrane repair (Couto *et al.*, 2017). Thus, our work ascribes a previously unacknowledged function for LAMP1 – targeting cytosolic proteins for lysosomal degradation. However, it remains unclear whether LAMP1 binds to PHD2 in lysosomes or in late endosomes (Geuze *et al.*, 1988). It remains to be determined how expansive is the repertoire of proteins that are degraded by this LAMP1-dependent pathway. It also isn’t clear how PHD2 interacts with LAMP1. Most likely the two proteins do not directly interact since the cytosolic tail is short and is required for its lysosomal targeting (Rohrer *et al.*, 1996). Thus, PHD2 likely interacts with LAMP1 via a complex composed of a chaperone and other proteins in a manner analogous to chaperone-mediated autophagy that uses hsc70 to facilitate cargo binding to LAMP2A (Cuervo *et al.*, 1994). For example, the cytosolic chaperone HSP90 can regulate lysosomal protein trafficking through a number of mechanisms including stabilizing lysosomal membrane protein complexes (Bandyopadhyay *et al.*, 2008, Liu *et al.*, 2009). PHD2 has been shown to bind to HSP90 and several co-chaperones (Song *et al.*, 2013) although this interaction promotes prolyl-hydroxylation of HIF-1α. However, it is possible that HSP90 complexed with other proteins can direct PHD2 to the lysosome and future work will identify the cytosolic complex required for LAMP1 dependent degradation of PHD2 and other substrates.

## MATERIAL AND METHODS

### Cells and Parasites

*To*x*oplasma gondii* type I RH strain tachyzoites were maintained by continuous passage in human foreskin fibroblasts (HFFs) with Dulbecco’s modified Eagle’s medium containing 10% heat-inactivated fetal bovine serum, 2 mM glutamine, penicillin/streptomycin. All other cells utilized in experiments (HeLa, HEK293T cells, and wild type or LAMP1KO MEFs) were also cultured in this medium. Intracellular parasites were collected from infected HFFs by scraping the infected monolayer and passing the material through a 27-gauge needle to liberate parasites from their host cells. All cells were tested for Mycoplasma contamination with the Mycoplasma Detection kit (Lonza; Basel, Switzerland) and found to be negative. Unless otherwise stated, reagents were purchased from Sigma (St. Louis, MO).

### Plasmids

FLAG-tagged PHD2 plasmid (p3XFLAG-PHD2) was provided by Dr. Richard Bruick (University of Texas Southwestern Medical Center). Human PHD2^Δ6-20^ and PHD2^Δ196-205^ were synthesized by IDT (Integrated DNA Technology Coralville, IA) with a-N-terminal 3XFLAG tag and cloned in pCMV-10 (Invitrogen, Carlsbad, CA). HA-VHL-pRc/CMv was purchased from Addgene (Cambridge, MA) (Plasmid #19999). pET24-GST-ODD (amino acids 402-603 in human HIF-1α) was synthesized by Genscript (Piscataway, NJ).

### VHL capture assay

Assays were done essentially as described in (Tuckerman *et al.*, 2004). Briefly, PNS extracts from mock- or parasite-infected cells were prepared as described above and then incubated in the presence of 1 mM α-ketoglutarate, 1 mM ascorbate and 50 µM FeCl_2_ for 90’ at 30°C with bacterially expressed GST-HIF-1αODD protein immobilized on glutathione sepharose beads. The beads were washed, resuspended in NETN buffer [20 mM Tris pH8.0, 100 mM NaCl, 1 mM EDTA, 0.5% NP40, 1 mM PMSF], and incubated with HA-tagged VHL synthesized using the TNT T7 Quick Coupled Rabbit Reticulocyte Lysate kit (Promega). The suspension was incubated overnight at 4°C with gentle rocking, analyzed by Western blotting, imaged using an Odyssey CLx infrared scanner (LI-COR, Lincoln, NE), and quantified with Image Studio 3.1 analysis software.

### Hydroxylation assays

Hydroxylation assays were either performed by matrix-assisted laser desorption/ionisation time-of-flight mass spectrometry (MALDI-TOF-MS) or by electrospray-ionisation liquid chromatography mass spectrometry (ESI-LC-MS). The following conditions were used: *Tg*PhyA *or Tg*PhyB (1 µM), *Hs*HIF1α CODD (DLDLEMLAPYIPMDDDFQL-NH_2_, 100 µM), *Hs*HIF1α NODD (DALTLLAPAAGDTIISLDF-NH_2_, 100 µM), or *Tg*Skp1^fl^ substrate (100 µM), (NH_4_)_2_Fe(II)(SO_4_)_2_ (50 µM), sodium L-ascorbate (1 mM) and 2-oxoglutarate disodium salt (500 µM) in HEPES (50 mM), pH 7.5. The reactions were incubated at 37 °C for 1 h and quenched with formic acid (1 % v/v). Peptide substrates were analysed by MALDI-TOF-MS using a Waters® Micromass® MALDI micro MX™ mass spectrometer, and protein substrates were analysed by ESI-LC-MS using a Waters® ACQUITY Xevo G2-S QToF mass spectrometer. Hydroxylation levels were quantified using MassLynx™ V4.0.

### Cell Fractionation Protocols

#### Nuclear/cytoplasmic fractions

Cells washed in cold-PBS were collected and homogenized using a Dounce homogenizer in a hypotonic buffer (20 mM Tris pH 8.0, 5 mM KCl, 1.5 mM MgCl_2_, 0.25mM DTT, 1mM PMSF, and 1X Protease Inhibitor cocktail (Sigma St. Louis, MO)). Lysates were centrifuged for 10’ at 10,000 x*g* at 4°C to separate cytoplasmic (supernatant) from nuclear fractions (pellet). Fractions were collected and analyzed by Western-blot.

#### S100/P100 Preparations

Cells were lysed in isotonic lysis buffer (0.3 M sucrose, 1 mM EDTA, 5 mM KCl, 120 mM NaCl, 20 mM Hepes pH 7.5, and 1X Protease Inhibitor cocktail) by passage through a 27 gauge needle. The lysates were centrifuged at 900 x*g* for 10’ (4°C) to pellet cell debris and then post-nuclear supernatants (PNS) were centrifuged at 100,000 x*g* for 1h at 4°C in a TLA120.2 rotor to obtain S100 (supernatant) and P100 (pellet) fractions. Carbonate extraction was performed by resuspending P100 fractions in isotonic lysis buffer containing 100 mM Na_2_CO_3_ pH 11.5 for 30’ at 4°C followed by an ultracentrifugation at 200,000 x*g* for 1h (4°C) in a TLA120.2 rotor. Protease protection assays were performed by incubating P100 fractions 15’ at 4 °C with 40µg/mL (final concentration) of proteinase K (Invitrogen) or the absence or presence of 1% Triton X-100. The reaction was stopped by adding 20mM PMSF, after which the sample was placed on ice for 10 min. The reactions were collected and analyzed by Western-blotting.

#### Density Gradient Fractionation

PNS in isotonic lysis buffer was adjusted to 40% Percoll and then centrifuged for 60’ at 34,000 x*g* in a TLA 110 rotor at 4°C. Alternatively, PNS was adjusted to 35% Optiprep and loaded at the bottom of a tube and layered with equal volumes of 25%, 20%, 15%, and 10% Optiprep in isotonic lysis buffer. Samples were centrifuged for 2 h at 200,000 x*g* in a TLS55 rotor at 4°C.

### Immunoprecipitation

Cells were collected and homogenized using a Dounce homogenizer in a Triton lysis buffer (50 mM Tris pH8.0, 150 mM NaCl, 1% Triton X-100, 10 mM NaF, 2 mM Na_3_VO_4_, 1 mM EDTA and 1X Protease Inhibitors). Lysates were centrifuged for 10 min at 10,000 x*g* at 4°C and 100µg of each lysate were incubated overnight at 4°C with indicated antibodies. Protein A-agarose beads (Cell Signaling Technologies; Danvers, MA) were added for 2 hours and after extensive washing the beads were analyzed by Western-blot.

### Luciferase assays

Cells were transfected using Lipofectamine 2000 (Invitrogen) as described (Spear *et al.*, 2006). Briefly 2×10^5^ cells were transfected with a total of 1 µg DNA consisting of PHD2 expression vectors (or empty vector control), firefly luciferase reporters (pHRE-Luc or pEGR4x-Luc) and pTK-Rel (Promega; Madison, WI). After 24 h, the transfected cells were mock- or *To*x*oplasma*-infected for 18 h and then analyzed using the Dual-Glo Luciferase Assay kit (Promega) according to the manufacturer’s protocol.

### siRNA Assays

As described (Menendez *et al.*, 2015), cells were transfected using RNAiMAX reagent (Invitrogen) with 10 nM of ATG5 (Invitrogen #s s18158, s18159, s18160), GAPDH, or negative control siRNAs. After 48 h, transfected cells were infected with RH type I parasites (MOI~4) for 18 hours after which whole cell lysates were prepared and analyzed by Western-blot.

## ACKNOWLEDGEMENTS

We thank Drs. Jason Kay and Sergio Grinstein for supplying the LAMP1KO MEFs and members of the Blader laboratory and Dr. Chris West for helpful discussions.

## FIGURE LEGENDS

**Supplemental Figure S1. Substrate Selectivity of *Toxoplasma gondii* PhyA and PhyB as determined by electrospray-ionisation liquid chromatography mass spectrometry (ESI-LC-MS) and matrix-assisted laser desorption/ionisation time-of-flight mass spectrometry (MALDI-TOF-MS).** (A) ESI-LC-MS confirms the activity of *Tg*PhyA on the S-phase kinase-associated protein 1 (*Tg*Skp1). By contrast, *Tg*PhyB did not hydroxylate *Tg*Skp1 under the used conditions. (B) *Tg*PhyA and *Tg*PhyB do not accept either the *N*-terminal and *C*-terminal oxygen-dependent degradation domains (NODD and CODD, respectively) of HIF1α as a substrates as determined by MALDI-TOF-MS analysis. Hydroxylation assays were carried out under the following conditions: *Tg*PhyA *or Tg*PhyB (1 µM), *Hs*HIF1α CODD (DLDLEMLAPYIPMDDDFQL-NH_2_, 100 µM), *Hs*HIF1α NODD (DALTLLAPAAGDTIISLDF-NH_2_, 100 µM), or *Tg*Skp1^fl^ substrate (100 µM), (NH_4_)_2_Fe(II)(SO_4_)_2_ (50 µM), sodium L-ascorbate (1 mM) and 2-oxoglutarate disodium salt (500 µM) in HEPES (50 mM), pH 7.5. The reactions were incubated at 37 °C for 1 h and quenched with formic acid (1 % v/v), before being subjected to analysis by ESI-LC-MS and MALDI-TOF-MS.

**Supplemental Figure S2. Down Regulation of Epitope-Tagged PHD2 by *Toxoplasma* is Resistant to MG-132.** FLAG-PHD2 transfected MEFs were mock or parasite-infected for 18 h and treated with or without MG132 for the last 6 h. Whole cell lysates were prepared and probed to detect FLAG-PHD2, IκBα, and β-actin.

**Supplemental Table S1: List of Antibodies Used in Study**

